# Remote Palpation of the Human Brain Using Simultaneous MR Elastography and Diffusion Tensor Imaging

**DOI:** 10.1101/2025.06.20.660588

**Authors:** Kulam Najmudeen Magdoom, Alexandru V. Avram, Joelle E. Sarlls, Peter J. Basser

## Abstract

“Remote palpation” appears to be an oxymoron, but here we demonstrate a non-contacting MRI method to obtain mechanical stiffness parameters of the human brain solely by measuring deformations caused by the pumping action of the heart. Mechanical stiffness is an important tissue property that is highly sensitive to subtle changes in the tissue milieu; MR elastography (MRE) is among a handful of methods used to measure it, typically via an external driver/tamper that introduces mechanical waves into the tissue. Applying MRE in the brain is challenging due to the use of an external actuator/tamper and the mechanical anisotropy of brain tissue, which requires a 4^*th*^-order tensor to describe it. In this study, we use the intrinsic deformation of brain tissue caused by periodic cardiac pulsations to measure the 4^*th*^-order elasticity tensor throughout the brain while simultaneously estimating the 2^*nd*^-order diffusion tensor in each voxel throughout the cardiac cycle which we use as *a priori* information in the reconstruction of the elasticity tensor. While the DTI-derived mean diffusivity (MD) appears uniform throughout brain parenchyma, stiffness maps obtained at about 1 Hz (i.e., at the fundamental cardiac frequency) show that brain tissue is very soft within gray matter, and within white matter pathways, such as along the corpus callosum, corona radiata, etc. Generally, stiffness differences at internal tissue boundaries are expected to produce local stress concentration there, which may predispose tissues to damage, e.g., in traumatic brain injury (TBI). Therefore, our novel tamperless MRE method has the potential to not only identify such interfaces, but assess and follow changes in tissue stiffness there that might occur following injury.

## Introduction

Understanding structure-function relationships in the human brain remains a *desideratum* with profound scientific implications and wide-ranging potential clinical applications. While it is currently not feasible to image whole brain structures *in vivo* at micro/nanometer length and nanosecond time scales, it is possible to measure and map various physical properties, such as magnetic susceptibility, water diffusivity, viscosity, bulk and shear moduli, hydraulic permeability, electrical conductivity, etc., at a larger continuum or mesoscopic length scale using magnetic resonance imaging (MRI) methods. These physical parameters inform biophysical models describing transport processes in tissue, appearing in equations governing the transport of magnetization, mass, momentum, energy, and charge, which underlie and constrain a myriad of basic physiological phenomena and processes. These transport parameters, however, also have a double-life in radiology and the neurosciences, serving as valuable quantitative imaging biomarkers. The best such examples are mean apparent diffusion coefficient (mADC) [1] and other diffusion tensor-derived quantities like the fractional anisotropy [2].

Here we focus on the tandem measurement of two key physical quantities: the shear modulus, which quantifies the resistance of a material to shear deformations, affecting the transport of momentum; and the diffusivity, which quantifies how thermally induced random tissue water transport results in molecular mixing. The shear modulus has been reported to vary by orders of magnitude among tissue types compared to other properties, such as magnetization relaxation rates, (i.e., *T*_1_ and *T*_2_) and bulk moduli [3, 4]. Water diffusivity on the other hand does not have a large dynamic range in tissue, and does not directly contribute to the shear modulus (as Newtonian fluids like water cannot support static shear stresses) thus providing complementary information about the tissue milieu. However, both properties are sensitive to the structural alignment of tissue components, such as axons, neurofilaments, and microtubules, which are ubiquitous in the brain [5, 6], and are expected to lead to orientational biases or anisotropy in transport processes. Based upon effective medium theory one expects a shared directional bias between anisotropic stiffness and anisotropic diffusion properties in tissue [1].

In particular, effective medium theory predicts that the stiffness and diffusion tensors share the same principal axes (i.e., eigenvectors), but that their corresponding principal values (i.e., eigen-values) may not be simply related to one another as they arise from different physical processes: diffusivity - from transport of mass or mixing via thermal collisions, and shear modulus - from the transport of momentum via adjacent mechanical coupling [1]. The nature of the partial differential equations (PDE) describing these transport processes are also qualitatively different: the diffusion equation being parabolic and the wave equation being hyperbolic with distinct solutions [7], whose parameters have different units and whose equations may possess different internal boundary conditions. Nevertheless, the microstructure of the medium can cause similarities in their respective anisotropic behaviors. These transport parameters are sensitive to features of material structure, composition, and organization in different ways. Because of the complementary information they provide, it is desirable, if possible, to measure diffusivity and shear modulus together in the brain, and examine their respective features, as we do here.

Diffusion tensor imaging (DTI) [1, 8] and MR elastography (MRE) [9] are imaging modalities that measure and map the the diffusion tensor and tissue shear modulus, respectively, throughout an imaging volume. Both have been under continuous development since the early to mid-1990s. DTI has been extensively used in the brain to identify white matter pathways [10, 11], study brain development [12] and diagnose a host of diseases, such as stroke [13] and cancers [14] via the DTI-derived mean apparent diffusion coefficient (mADC) or mean diffusivity (MD) [15]. Meanwhile, MRE has been primarily used to assess liver fibrosis [16] with limited applications to characterize brain tissue due to several factors we describe below.

MR elastography, as originally proposed, measures displacements resulting from shear waves with a prescribed frequency applied to a sample using an external actuator or tamper. These displacements are then used to estimate the shear modulus based on a model that relates the material’s strain and stress [9]. Applying MRE in the brain is challenging for several reasons; 1) Brain tissue is mechanically anisotropic [17] which requires a 4^*th*^-order elasticity tensor with 21 unknowns to fully describe its mechanical properties, resulting is an ill-posed problem [18], and 2) the external tamper itself is costly and requires special training to operate. The tamper often operating between 25-60 Hz [19] may also be less sensitive to poroelastic drainage effects, which are thought to drive physiological glymphatic brain clearance mechanism [20] through the intrinsic pulsations of brain tissue around 1 Hz from the heart.

Mechanical anisotropy is largely ignored in many brain MRE studies owing to the difficulty of measuring the entire 4^*th*^-order tensor [21–24]. This omission, however, introduces additional variability in the measured isotropic tissue mechanical properties as these become dependent on the placement of the external tamper with respect to the brain position and orientation across subjects [25]. Several studies have attempted to quantify mechanical anisotropy with MRE by solving a highly ill-posed problem. The full 4^*th*^-order elasticity tensor has as many as 21 unknown elements but only 3 equations governing the material displacement are typically available.

A common approach is to assume additional underlying symmetries, such as transverse isotropy of the material in coherent fibrous regions, which reduces the number of unknowns from 21 to five. This number can be further reduced to three by assuming tissue incompressibility [26]. This simplification, however, requires *a priori* knowledge of the symmetry axis and hence typically limited to skeletal muscle tissues or phantoms with known principal axes [27–29]. To try to address this, Qin et al. had used principal directions from a separate DTI scan to estimate the transverse isotropic stiffness tensor in phantoms with a single fiber direction [30]. Recently, Smith et al. presented a non-linear reconstruction approach on live brain tissue with a tamper vibrated along two different orientations and solved for material parameters using a computationally intensive finite element method (FEM) model analysis framework with principal directions obtained from separate DTI scans [31]. The most general stiffness tensor estimated in the brain to date was obtained using waveguide elastography in which the three principal axes of the diffusion tensor were assumed to coincide with principal frame of the stiffness tensor, simplifying the form of the governing equations and the estimation of the orthotropic stiffness tensor (i.e., having 9 independent elements) [6]. The approach is nonetheless computationally intensive and it was implemented using displacements obtained using an external tamper.

Given the complexity and experimental requirements for conventional brain MRE using a tamper, a more pragmatic approach would be to perform brain MRE by employing the intrinsic or endogenous deformation of the brain tissue caused by the subject’s own cardiac pulsations, overcoming the aforementioned drawbacks of MRE using an external tamper or mechanical actuator. In this embodiment, the phase images encode the tissue motion to enable the estimation of the displacements, while the magnitude images, which are sensitive to molecular diffusion, provide a means to concurrently estimate the fiber orientation within a voxel, potentially to inform the estimate of the elasticity tensor. A hurdle to overcome, however, is that the resulting brain tissue deformation is still small for conventional imaging acquisitions (i.e., on the order of several microns within a 8 mm^3^ voxel) [32]. This however is a benefit for MRE as the stress-strain relationship can be assumed to be linear in this infinitesimal strain regime, simplifying stiffness reconstruction and mapping. It should be noted that the idea of simultaneously measuring both diffusivity and displacement/velocity dates back to the ‘50s by Carr and Purcell [33], and later by Stejskal [34] in the ‘60s and in the ‘80s by Callaghan et al. [35]. Recent applications involve simultaneous DTI and MRE in mouse [36] and human brain [37] using a tamper, resulting in large tissue deformations compared to the intrinsic pulsations making them easier to detect, and assuming the mechanical properties of the brain to be isotropic. Their method is also immune to phase errors compared to intrinsic MRE since motion encoding was performed using oscillating gradients (OGSE) tuned to the frequency of motion.

It should be noted that there have been several studies describing MR imaging of heart-driven brain displacements [32, 38–42] with attendant strengths and limitations. Some of these methods use stimulated echo displacement encoding (DENSE), which can be difficult to implement and are not SNR efficient (e.g., there is a 50% SNR loss using a stimulated echo (STE) sequence as compared to a spin echo (SE) sequence), while others use video amplification of magnitude images acquired over the cardiac phase. Weaver et al., and Ingeberg et al., inverted these displacements into isotropic shear moduli using a computationally intensive FEM framework reporting orders of magnitude differences in their estimated stiffness values (8 kPa vs 200 Pa, respectively) [23, 43]. Herthum et al. estimated stiffness values of tens of Pascals [44] based on the wave speed measured across a brain slice. Given the large inconsistency among these experiments and the importance of measuring mechanical properties of brain tissue, there is still a need to develop a robust and accurate method to measure stiffness tensor throughout the brain and across subjects using only the intrinsic pulsations produced by the beating heart.

In this study, we introduce a new experimental design to reliably acquire whole-brain complex-valued 3D motion-encoded MRI data throughout the cardiac cycle, by using standard pulsed gradient spin echo (PGSE) MRI measurements. We use an outlier rejection strategy to overcome spurious phase errors and build up consistent complex-valued 3D motion-encoded MRI data volumes throughout the cardiac cycle. We estimate the diffusion tensor field from the magnitude of the MRI signal and compare its features with the simultaneously measured displacement vector field throughout the cardiac cycle. We concomitantly reconstruct the elasticity tensor in the whole brain from the measured displacement field (obtained from the phase signal), assuming transverse isotropy in white matter, where the symmetry axis of the elasticity tensor is expected to be given by the principal eigenvector of the estimated diffusion tensor using the “cross-property” concept [1]. Finally, we determine the form of the elasticity tensor in each voxel by inverting the momentum conservation equation.

## Materials and Methods

### MRI acquisition and processing

MRI data was acquired in six healthy volunteers who provided informed written consent to participate in the NINDS IRB approved imaging research protocol on a 3T scanner (Prisma, Siemens Healthineers) with 80 mT/m peak gradient strength and a 200 T/m/s slew rate using 20-channel receive RF coil. Whole-brain data were acquired using standard PGSE echo planar imaging (PGSE-EPI) sequence with motion-encoding gradients (MEG) played along the six directions of the icosahedron [9] at b = 350 *s/mm*^2^ and *v*_*enc*_ = 0.7 mm/s along with a b = 0 s/*mm*^2^ scan using the following parameters: *δ/*Δ = 7/48 ms, field of view = 210 × 210 × 120 mm, GRAPPA acceleration factor = 2, TR/TE = 5,600/71 ms, NEX = 144 per direction and slice, and a 2 mm isotropic spatial resolution. The total scan time for this study was approximately 1.5 hours, long enough to obtain at least two sets of full displacement encoded volume (i.e., all directions and slices) with a 100 ms bin size for test-retest analysis. The MEG pulses were kept narrow with long time interval between them to enhance scan sensitivity to coherent brain pulsations while reducing signal loss from diffusion [45]. The pulse-oximeter signal and MRI triggers were simultaneously recorded using a Biopac System (Biopac, Goleta, CA, USA) for retrospective gating.

The MRI phase was unwrapped using a Fourier-based method [46] after removing static phase offsets by complex division by the *b* = 0 *s/mm*^2^ scan. The linear phase errors arising from eddy currents, rigid body motion, etc., were removed using linear regression. The *B*_0_-induced geometric distortion was corrected using FSL’s Topup software [47, 48]. The displacement-encoded images were then partitioned into ten different bins each approximately 100 ms long covering the entire cardiac cycle using the measured pulse-oximeter (SpO_2_) waveform to indicate phase in the cardiac cycle. It should be noted that the number of bins for a given dataset depends on the minimum number of repetitions required for the full displacement encoded volume (i.e., all directions and slices), higher number of bins can be achieved by reducing the number of repetitions per bin and vice-versa but ultimately is limited by subject’s heart rate and TR per slice.

### Outlier rejection and displacement estimation

MRE reconstruction involves computing both spatial and temporal derivatives, which require smoothly varying 4D displacement fields (i.e., three dimensions in space and one dimension in time or frequency). The displacement field is however not measured instantaneously since it requires data from multiple directions, which are acquired over multiple scan repetitions. Given the high scan sensitivity to bulk motion and the multitude of factors affecting the signal phase, complex phase errors are introduced between these acquisitions in live brain MR imaging data, especially along the slice-encoding direction in EPI. These errors will not be corrected by linear regression and can arise from brain sloshing accompanying unpredictable head motion [32], phase errors from heating of passive shims [49], etc. While previous work by Barnhill, et al. addressed this issue by using dejittering algorithms and wavelet transforms to remove these errors [50], we employ a simpler method to reject the inconsistent phase measurements in a voxel from the multiple repetitions of data acquired in each bin for a given motion-encoded gradient (MEG) direction. Assuming a non-zero signal magnitude, the voxelwise phase distribution over multiple repetitions follow a Gaussian profile. Therefore, for each MEG direction, bin and voxel, we exclude from the estimate of MRE and DTI data specific repetitions whose phase values fall outside one standard deviation of the mean phase. It should be noted that the number of repetitions which were eventually averaged for a given slice varies depending on the slice location and MEG direction.

Given the linear relationship between phase and displacement, the consistent phase images from all MEG directions are then Fourier transformed in time and converted to harmonic displacement vectors using linear regression [51]. The measured displacement field is then spatially smoothed using a local 3D Gaussian kernel prior to calculating spatial derivatives required for the stiffness estimation.

### Diffusion tensor estimation

The magnitude signal encodes information about the diffusion of water molecules. We acquire a sufficient number of DWI measurements to simultaneously estimate the diffusion tensor field within each cardiac bin [1, 8]. Since the image phase is more sensitive to motion than magnitude, the magnitude of those repetitions with inconsistent phase were automatically rejected. The consistent magnitude signals were averaged over each bin and direction, and denoised using Marchenko-Pastur principal component analysis (MP-PCA) [52] implemented in the MRTrix software environment [53] to estimate the time-varying diffusion tensor field using non-linear least squares approach with a positive definiteness constraint applied on the diffusion tensor, and positivity constraint applied on *S*_0_ to ensure physicality. The various diffusion tensor-derived parameters, such as fractional anisotropy (FA) a.k.a. diffusion anisotropy, MD, direction-encoded color (DEC) maps and diffusion tensor-orientation distribution function (DT-ODF) were all calculated from the estimated diffusion tensor at each phase in the cardiac cycle [2, 54, 55].

### Helmholtz decomposition of displacement field

The measured 3D displacement field is decomposed into its longitudinal (**u**_*L*_) and transverse (**u**_*T*_) components using the Helmholtz decomposition to investigate the spectral features of the longitudinal and shear wave fields, respectively [56]. These components are written using the scalar potential field, (Φ), and solenoidal vector potential field, (**A**), respectively, as shown below,

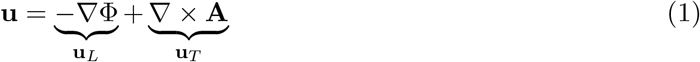

The individual components are estimated by solving the Poisson equations below obtained by taking the divergence and curl of the above equation,

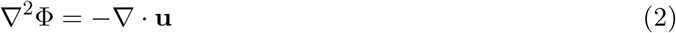

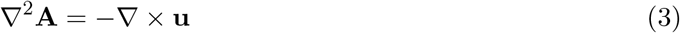

The above equations are solved for the scalar and vector potentials using an algebraic multigrid method [57] with their values set to zero outside the brain, which was segmented from the imaging volume using the brain extraction tool (BET) implemented in FSL [58, 59].

### Governing equation for the displacement field

The equation governing the evolution of the displacement field of an elastic medium in time and space is the momentum conservation principle (i.e., Navier’s equation),

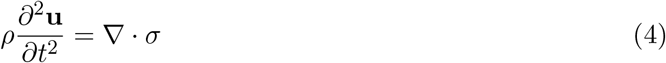

where *σ* is the 2^*nd*^-order stress tensor and *ρ* is the density of the medium. Assuming a general linearly elastic (i.e., Hookean) material, *σ* = C : *ε*, where C is the 4^*th*^-order elasticity tensor and *ε* is the 2^*nd*^-order strain tensor, which is given below,

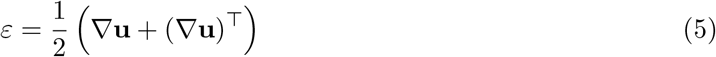

Applying transverse isotropic symmetry about the *z*-axis, the 4^*th*^-order elasticity tensor can be expressed as a 6 × 6 matrix in the principal frame with five unknowns as follows [18],

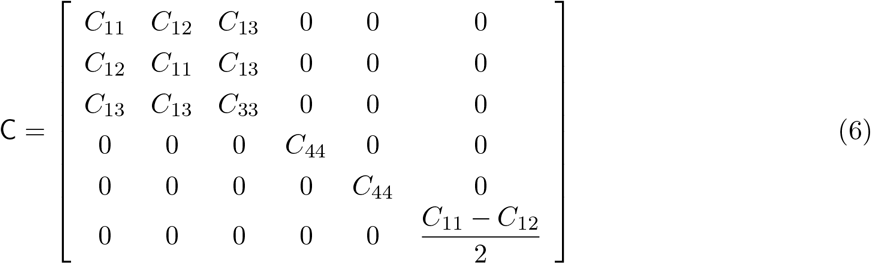

where *C*_*ij*_ are the various moduli that characterize the tensor. Substituting Equations (5) and (6) into Equation (4) results in the following equations for the displacement,

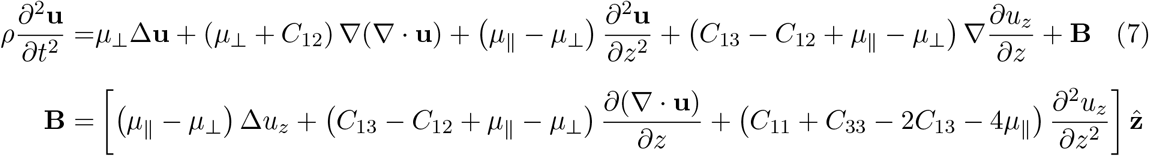

Where 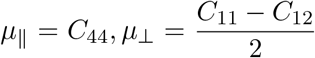 are the shear modulus parallel and perpendicular to the fibers, respectively, 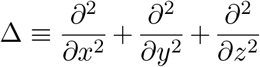 is the Laplacian operator, and 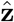 is the unit vector along *z*. The above set of equations provide a complete description of wave propagation in a transverse isotropic linearly elastic medium.

### Estimation of the elasticity tensor

#### Isotropic tensor model

In gray matter, we assume an isotropic stiffness tensor. Practically these are voxels whose FA in the quiescent phase of the cardiac cycle is less than 0.2. For an isotropic stiffness tensor, the following relations apply: 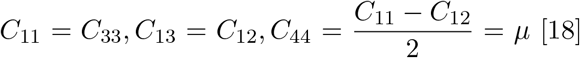 [18]. Substituting these in Equation (7), and performing a Fourier Transform with respect to time, *t*, the governing equation for the time-harmonic displacement field, **û**, is given by,

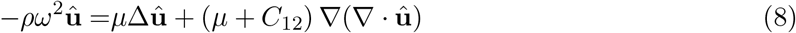

where *ω* is the angular frequency equal to 2*πf* with *f* being the fundamental frequency of heart beat. The two unknown parameters are computed by rewriting the above governing equations as an optimization problem that is solved in each voxel,

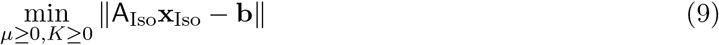

Where *µ* is the shear modulus, 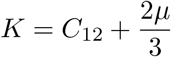 is the bulk modulus, A_Iso_ is a 3 × 2 complex valued matrix representing the operator multiplying the vector of unknown parameters, **x**_Iso_, which in this case consists of *C*_11_ and *C*_12_, and **b** is the vector of terms on the left hand side of the governing equations (i.e., Equation (8)). The positivity constraint was applied on the shear modulus and bulk modulus to ensure physicality. The optimization problem was solved using the MOSEK solver [60] in the CVXPY software environment [61, 62] with displacement derivatives estimated using the Savitzky-Golay filter [63].

#### Anisotropic stiffness tensor model

In regions with large FA (i.e., FA *>* 0.2), which we assume to be along white matter pathways, we use a transverse isotropic model. An *Ansatz* we also apply is that both the elasticity and diffusion tensors are assumed to share the same principal coordinate frame [1, 6]. The five unknown tensor elements are estimated by solving the following optimization problem using the three equations of motion similar to the isotropic model (i.e., Equation (7)),

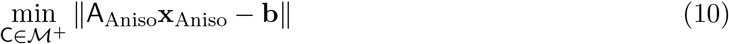

where ℳ ^+^ is the manifold of positive definite matrices, A_Aniso_ is a 3 × 5 complex valued matrix representing the operator multiplying the vector of unknowns of the elasticity tensor, **x**_Aniso_. The positivity constraint for the isotropic shear and bulk modulus translates to a positive definiteness constraint for the elasticity tensor to ensure physicality [64]. It should be noted that since the displacement field is complex, we have six equations and six constraints from applying Sylvester’s criterion for positive definiteness to solve for five parameters thereby guaranteeing a well-posed problem.

Since the above equations assume the axonal fibers are oriented along the z-axis, the displacement field and its derivatives are rotated appropriately in each voxel using a rotation matrix, R, derived from the principal eigenvector of the corresponding diffusion tensor in the quiescent phase of the cardiac cycle using the following relations in tensor notation,

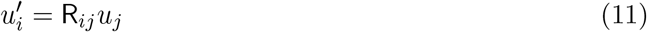

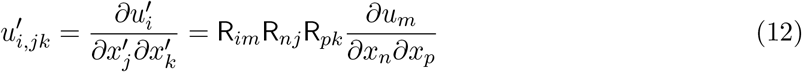

The estimated 4^*th*^-order stiffness tensor is then rotated into the lab coordinate frame using the inverse rotation matrix (i.e., its transpose, R^⊤^) to map its features across voxels and correlate its components with the diffusion tensor, as shown below,

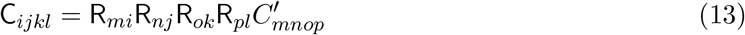

N.B., we ensure that R is a proper right-handed rotation matrix.

### Visualization

The kinematics of the motion is visualized using a 3D vector plot of the measured harmonic displacement field amplitude and its curl along with a scalar map of the divergence. The diffusion tensor derived maps are shown as a function of the phase in the cardiac cycle to investigate their variability due to brain motion and/or deformations. The DEC maps are warped using the amplified displacement field measured in each cardiac phase to delineate the extent of deformation on various white matter tracts using the PyVista software environment [65]. Streamline tractography is performed using the measured DT-ODF in the MRTrix software environment [53], and the tracts are colored according to the value of quantities we obtain from the estimated diffusion and elasticity tensor, as discussed below.

Features derived from the estimated elasticity tensor are visualized using a set of “stains” and glyphs. The mechanical anisotropy, MA, is a measure of the deviation of the elasticity tensor from its isotropic equivalent, C_*iso*_, as shown below adapted from [66],

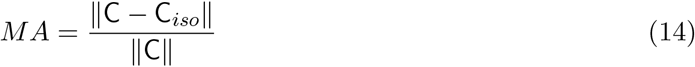

It should also be noted that the definition of FA for diffusion anisotropy is analogous to the MA. The isotropic part of an elasticity tensor is given by averaging it over all orientations resulting in the following [67],

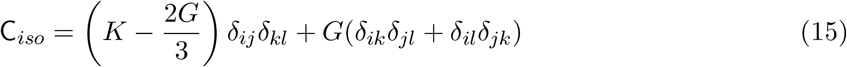

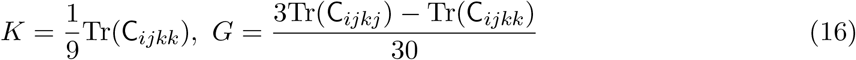

where *δ* is the Kronecker delta, and *K* and *G* are the effective isotropic bulk and shear moduli, respectively, for a given elasticity tensor. N.B., the Einstein convention is used, meaning that repeated indices imply summation over that index. While the true bulk modulus of tissue is difficult to estimate given the wavelength associated with sound waves in water at 1 Hz are orders of magnitude larger than the brain size, this becomes an “apparent” bulk modulus reflecting the squeezing of fluid in and out of the voxel during the long actuation period (i.e., poroelastic effect). Shear modulus should also be interpreted as an “apparent” shear modulus given we are not separately accounting for the solid-fluid coupling that could occur at the low actuation frequency. The shear modulus is also expressed as a function of local wavelength, *λ*, to account for heart rate variations across subjects which is given by,

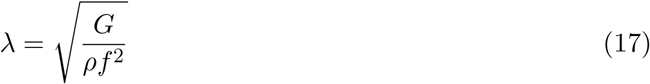

We can map 4^*th*^-order tensors to glyphs by contracting them with unit radial vectors. The orientation dependent axial stiffness, *E*(**n**), and shear stiffness, *G*(**n, m**), is given by the following expressions,

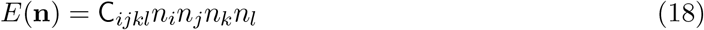

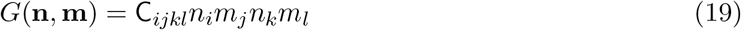

where **n** is the unit vector on a sphere, and **m** is an unit vector perpendicular to **n**. The shear stiffness for a given **n** averaged over all lateral orientations, **m**, is obtained using the identity, 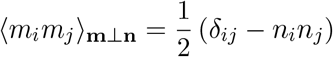, proven in the Appendix,

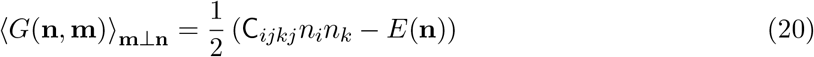

It can be observed that the above relation reduces to the shear modulus for the case of isotropic elasticity tensor. The spherical functions representing the axial stiffness and orientationally averaged shear stiffness are displayed as 3D glyphs.

## Results

First, we examine features of the displacement vector field throughout the cardiac cycle. Specifically, we display the real and imaginary parts of the displacement field amplitude oscillating at the cardiac frequency, which is approximately 1 Hz, in Figure 1 along with the animation of the displacement vectors in the time domain (in Animation 1 in supplementary material). An effective means to visualize these displacement vector fields is to superimpose them onto the DTI-derived FA map obtained at the quiescent phase of the cardiac cycle. The FA map provides an anatomical reference frame to help locate brain structures whose displacement vector field glyphs are also displayed. The displacement at the cardiac frequency peaks at around 150 *µ*m in the brainstem in the superior-inferior direction corresponding to a velocity of approximately 1 mm/s. The L-R gradient in the measured displacement field suggests a funnel shaped deformation of the brain with the brain stem and spinal cord acting like a piston. The imaginary part of the harmonic displacement field was comparable to the real part at the base of the brain and decays faster than the real part in the direction of the head.

**Figure 1:**
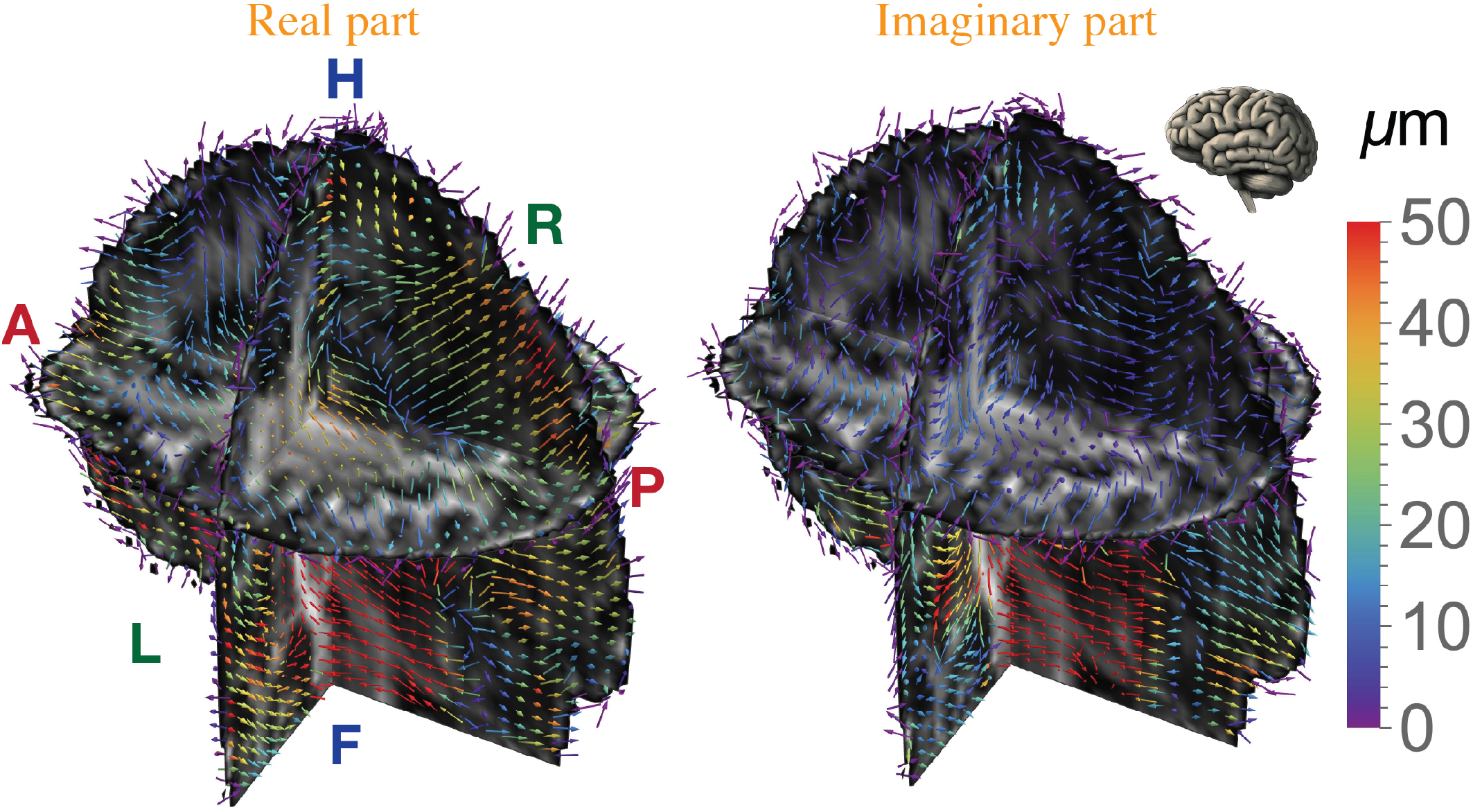
3D vector plots of the real and imaginary parts of the displacement field oscillating at the cardiac frequency (i.e., ≈ 1 Hz) for a representative healthy volunteer. The fractional anisotropy (FA) map obtained at the quiescent phase of the cardiac cycle is overlaid on the displacement field to provide anatomical context. The displacement vectors are colored based on their magnitude; the anatomical axes (A - Anterior, P - Posterior, H - Head, F - Feet, R - Right, L - Left) are indicated. The increasing F-H component of the displacement field from medial to lateral portions of the brain suggests a funnel-shaped profile along the F - H axis.

Second, we explore the repeatability or reproducibility of the displacement vector field measurements from which the elasticity tensor is computed using a test/retest paradigm. This is possible given that we can perform more scan repetitions in the same imaging session than is required to acquire a single tranche of MRE data. The harmonic displacement vector field obtained at the cardiac frequency in two repetitions are shown in Figure 2. We see high reproducibility with data obtained during different times in the scanning session on the same volunteer. The displacement profiles are noisier than the displacement vector field shown in Figure 1 but features such as the displacement magnitude and shape of the profile are highly reproducible.

**Figure 2:**
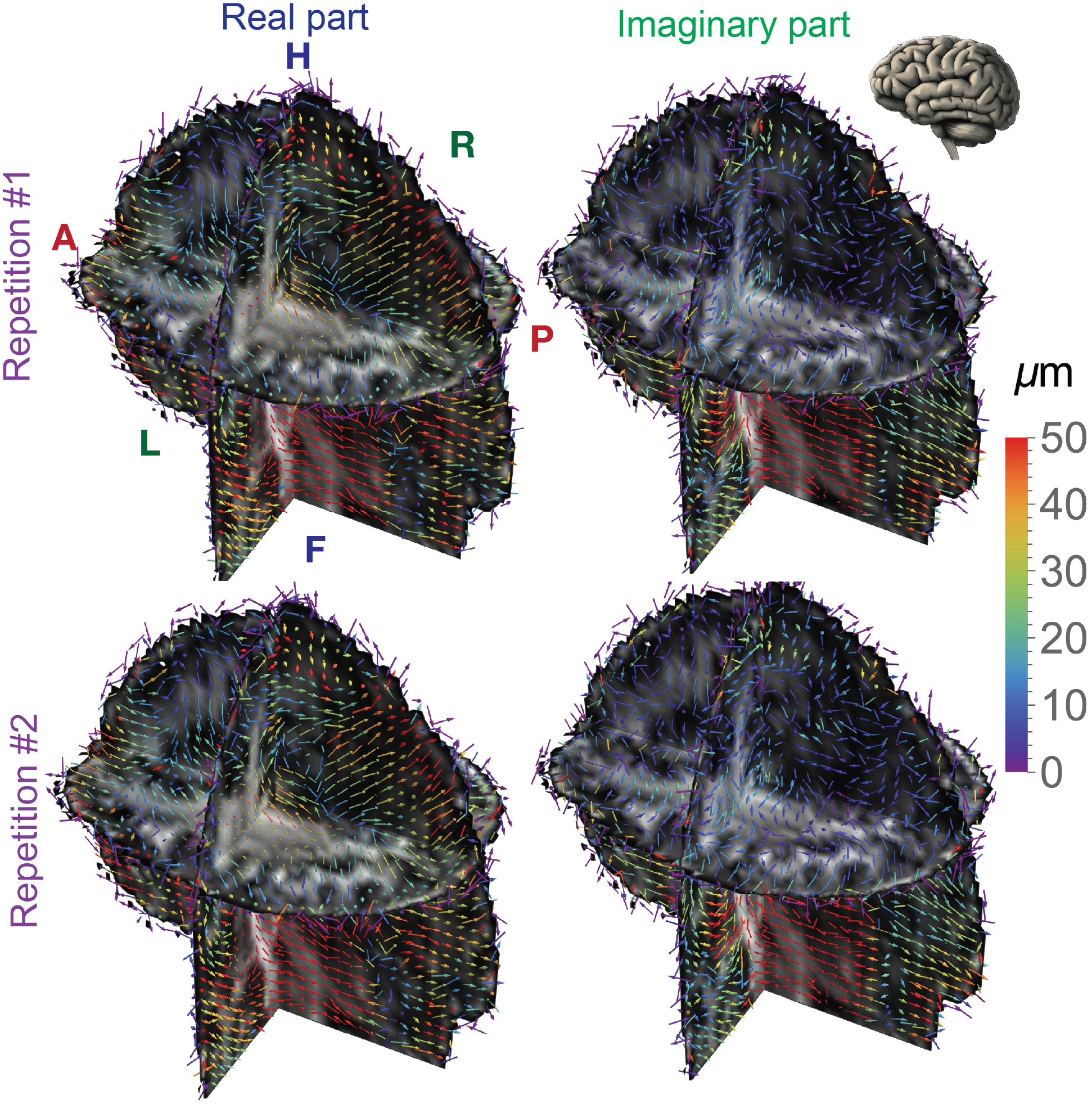
Test-retest repeatability of the displacement field measurement shown using 3D vector plots of the real and imaginary parts of the displacement field oscillating at the cardiac frequency (i.e., ≈ 1 Hz) for a representative healthy volunteer. The fractional anisotropy (FA) map obtained at the quiescent phase of the cardiac cycle is overlaid on the displacement field to provide anatomical context. The displacement vectors are colored based on their magnitude, and anatomical axes (A - Anterior, P - Posterior, H - Head, F - Feet, R - Right, L - Left) are indicated. The displacement fields were highly reproducible albeit noisier than the mean displacement vector field.

Third, we show the complex valued compressive and shear strains in the brain tissue resulting from cardiac pulsations using maps of divergence and curl of the harmonic displacement field overlaid on the FA map and expressed as percentage in Figure 3. Overall, the curl is higher in magnitude than the divergence but both have a median less than 0.2% over the entire brain. By comparison with the FA map, the curl and divergence are similarly heterogeneous in gray and white matter regions.

**Figure 3:**
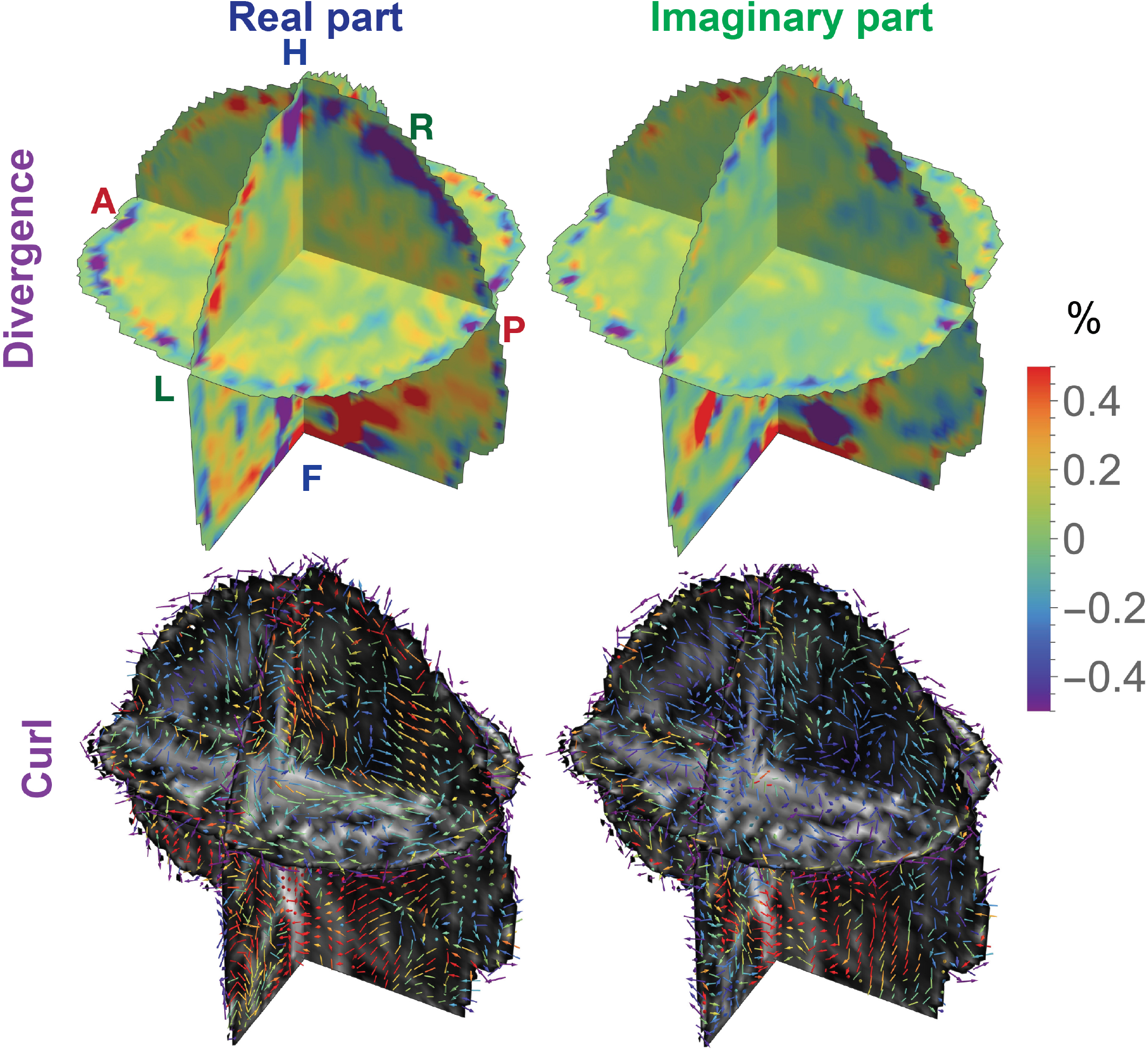
The real and imaginary parts of the divergence and curl of the harmonic displacement fields shown in a “3-D slicer” format for a representative healthy volunteer. The dilatation is a scalar map showing volumetric changes, depicted using 3D vectors overlaid on the fractional anisotropy (FA) map to provide anatomical context. The curl vectors are colored based on their magnitude; their anatomical axes (A - Anterior, P - Posterior, H - Head, F - Feet, R - Right, L - Left) are indicated. The divergence in the brain tissue was very small but above the noise level with the curl having an overall higher magnitude. The gradients in curl field were noted in the brain parenchyma suggesting deformation.

We show the complex valued longitudinal and transverse wave fields in the brain at the cardiac frequency along with the results of wave number analysis in Figure 4. Both the wave fields are non-zero and exhibit complex contrast in the brain tissue. The wave number analysis showed a continuum of wavelengths at the low cardiac frequency with a characteristic bump in some transverse and longitudinal components beginning around 0.1 *mm*^−1^, corresponding to a shear modulus of 0.1 Pa.

**Figure 4:**
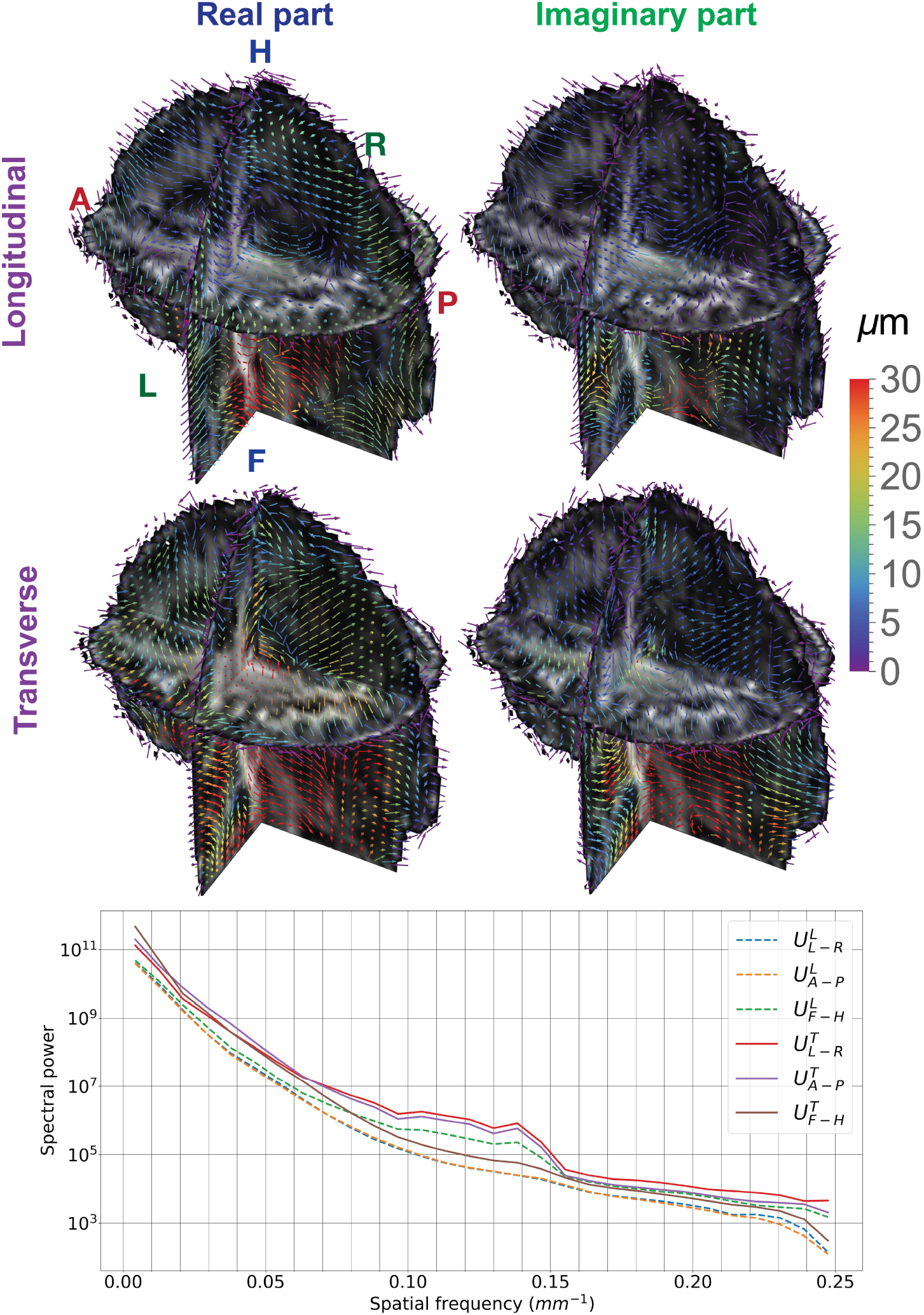
Wave number analysis of transverse and longitudinal components of the intrinsic 3D displacement field measured in the whole brain from a representative subject. A Helmholtz de-composition of the complex-valued harmonic displacement field oscillating at the cardiac frequency separates the longitudinal and transverse components, which are shown using 3D vector plots. The three components of the longitudinal and transverse wave vector fields were Fourier transformed and the resulting 3D k-space data was radially segmented into 30 bins whose spectral power in each bin is plotted as a function of the associated wave number shown in the bottom row. The presence of a range of wavelengths resulting from mechanical dispersion is clearly visible in these plots especially the hump around 0.1*mm*^−1^, which corresponds to a wavelength of 10 mm that results in a shear modulus of roughly 0.1 Pa. (A - Anterior, P - Posterior, H - Head, F - Feet, R - Right, L - Left)

We report the diffusion tensor and displacement vector fields at different phases of the cardiac cycle in a representative axial slice from a healthy volunteer in Figure 5 where we observe variations in the diffusion tensor field as a function of the cardiac phase. We also show the DEC map in the three orthogonal planes, which are deformed by the measured displacement field amplified 40x at various phases of the cardiac cycle (see Animation 2 in supplementary material) that show the deformation of various fiber tracts. The MD map showed variations as high as 45% in the corpus callosum (red arrows in the figure), and DEC maps showed variations in conspicuity of certain white matter fiber tracts (white arrows in the figure), which correlated with the sign of the F-H component of the displacement vector. For example, the MD was higher in the corpus callosum (i.e., red arrow) when the F-H component of the displacement vector was positive and vice versa due to the displacement of the ventricular CSF underneath it. The animation shows deformation in all parts of the brain.

**Figure 5:**
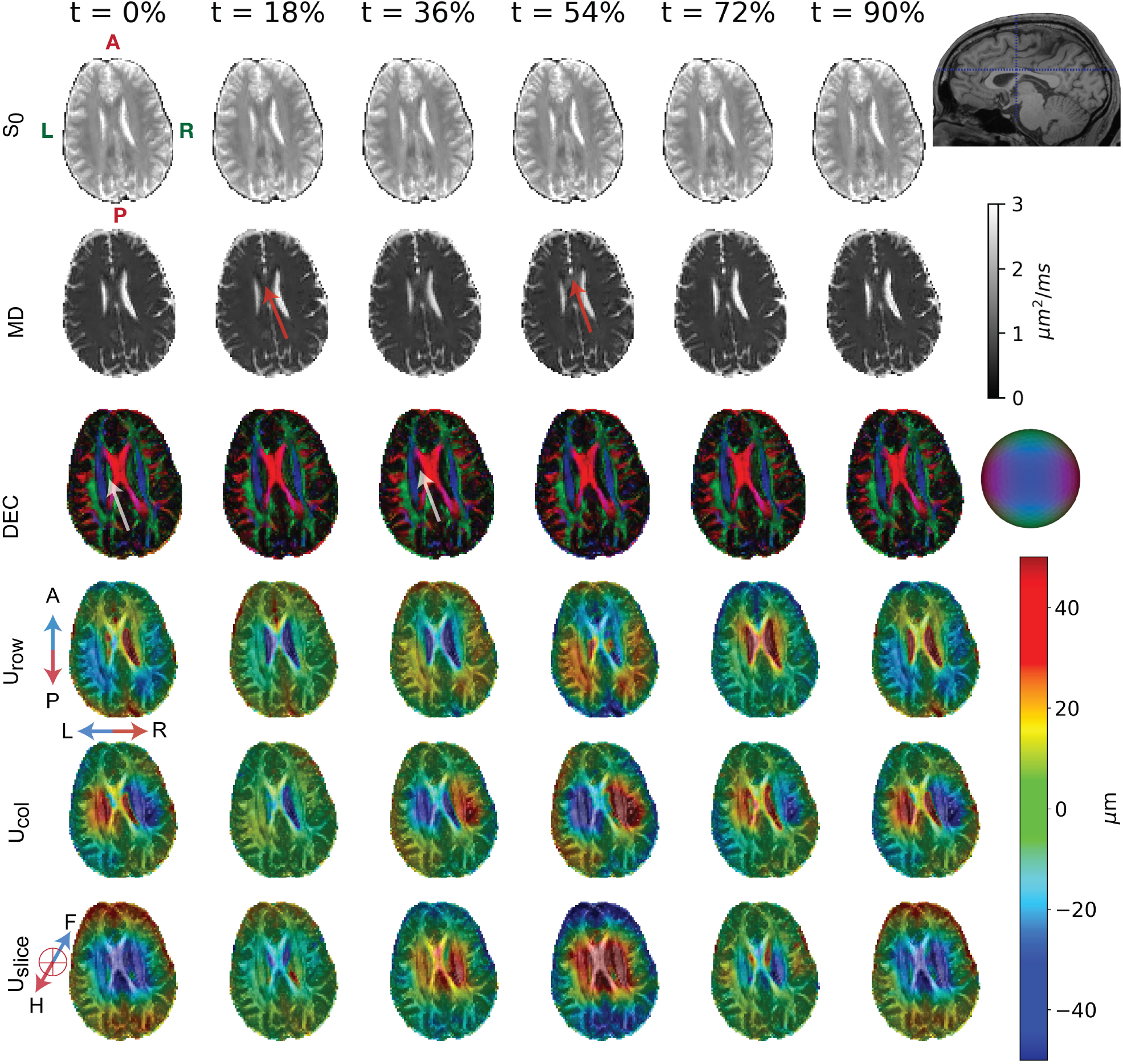
Time-varying diffusion tensor and displacement vector fields at different phases of the cardiac cycle for a mid-brain axial slice from a representative healthy volunteer. The DTI-derived maps including *S*_0_, mean diffusivity (MD), and direction-encoded color (DEC) along with the color sphere legend are displayed as a function of the cardiac phase (given as percent of total period after the *p*SO_2_ trigger). The three components of the displacement vector field in the patient coordinate system (A - Anterior, P - Posterior, H - Head, F - Feet, R - Right, L - Left) are provided for comparison with a map of fractional anisotropy (FA) overlaid for anatomical context. The location of the slice in the sagittal plane of an anatomical scan is provided on the top right using a violet dashed line. For clarity, the time axis is down-sampled in the figure by a factor of 2. Some of the variations in the DTI “stains” across the cardiac cycle are likely due to partial volume effects induced by pulsations, are shown using arrows. White arrows represent the variations in FA in CSF-filled ventricles across the cardiac cycle, and the red arrows represents the increase in MD in the corpus callosum.

We compare the diffusion and elasticity tensor maps in axial, coronal, and sagittal slices from a representative volunteer in Figure 6 using a 3D slicer representation and performed region of interest (ROI) analysis across subjects which are tabulated in Table 1. The diffusion tensor maps include MD and FA while the elasticity tensor maps include the isotropic shear, bulk modulus, and mechanical anisotropy. The MD is fairly uniform in brain parenchyma (≈ 0.7*µm*^2^*/ms*) while the shear and bulk modulus maps were highly heterogeneous. White matter overall had higher stiffness compared to gray matter. Both isotropic shear and bulk modulus were small at the cardiac frequency with an average approximately 0.67 Pa and 0.46 Pa, respectively for the entire brain across subjects as shown in the table. The mechanical anisotropy in white matter is roughly 60% higher compared to diffusion anisotropy. The coefficient of variation in shear and bulk modulus is an order of magnitude larger than for the mean diffusivity. Differences in shear modulus were observed among various brain regions, for example, the internal capsule was stiffer than the corpus callosum observed across multiple subjects. Even though the diffusion tensor eigenvectors were used to estimate the stiffness tensor, the extent of mechanical anisotropy was different from that of FA. The mechanical anisotropy of several areas of the brain were accentuated compared to FA.

**Table 1:**
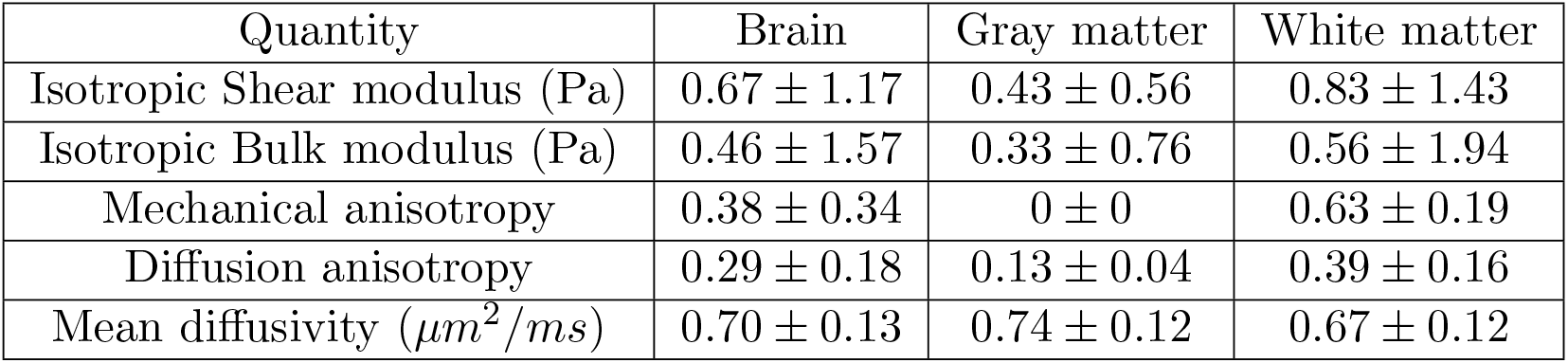
Region of interest (ROI) mechanical and diffusion property values at cardiac frequency averaged over all six subjects. Gray and white matter were segmented based on their fractional anisotropy and mean diffusivity values. The brain is composed of both gray and white matter. White matter overall had higher stiffness compared to gray matter. Both isotropic shear and bulk modulus were small at the cardiac frequency. The mechanical anisotropy in white matter is roughly 1.5x the diffusion anisotropy. It should be noted that the zero mechanical anisotropy for gray matter is by design. The coefficient of variation in shear and bulk modulus is an order of magnitude larger than mean diffusivity.

**Figure 6:**
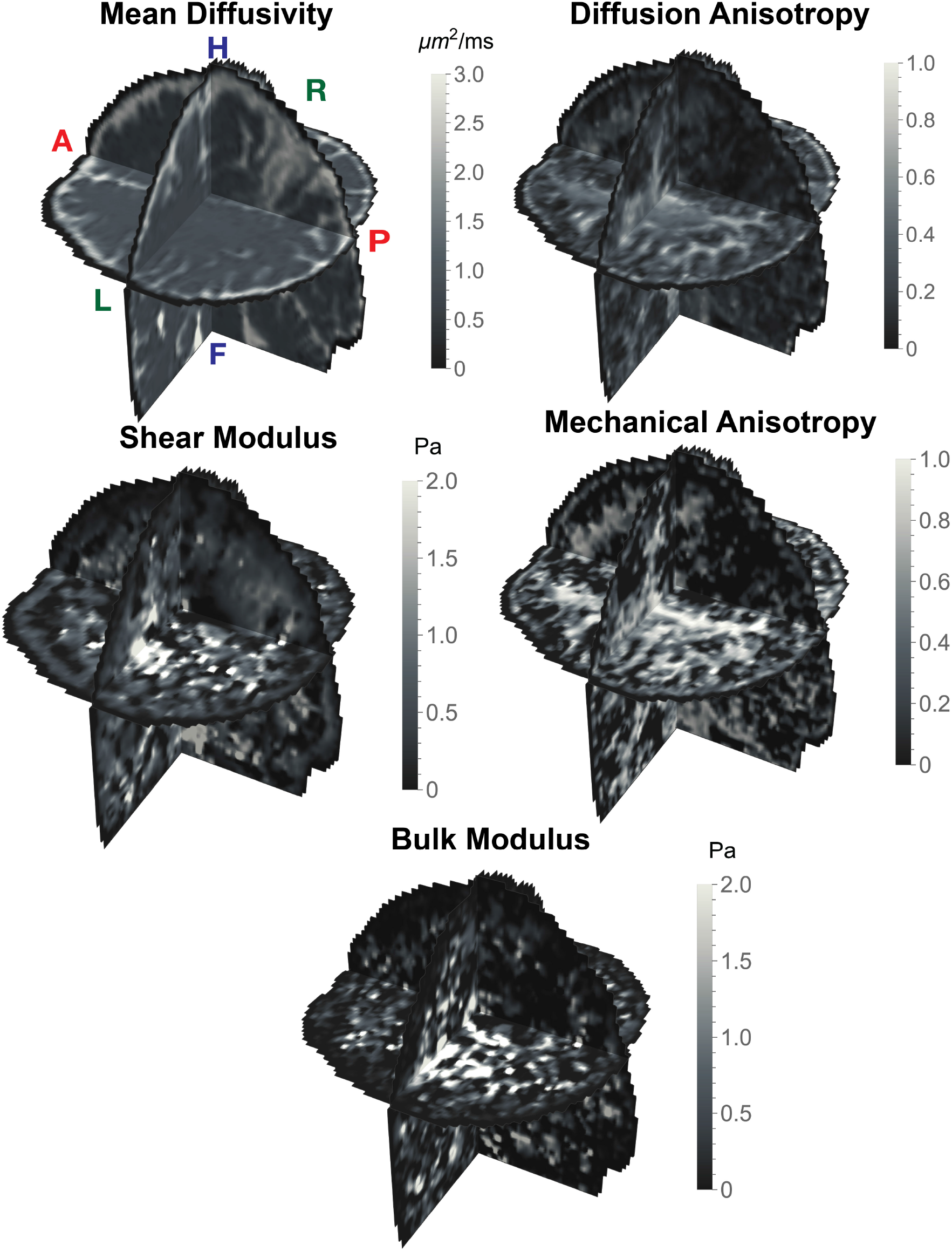
Diffusion and elasticity tensor maps along axial, coronal, and sagittal slices from a representative healthy volunteer. These include the mean diffusivity (MD), Fractional Anisotropy (FA), mechanical anisotropy, isotropic shear modulus and bulk modulus. The anatomical axes (A - Anterior, P - Posterior, H - Head, F - Foot, R - Right, L - Left) are indicated in the figure. White matter is stiffer than gray matter with differences noted among various white matter regions. The bulk and shear modulus were heterogeneous with gray-white matter contrast while MD was uniform. The mechanical anisotropy was accentuated compared to diffusion anisotropy.

We investigated the 2D distribution of diffusivity and mechanical properties in gray and white matter regions of the whole brain, and possible correlations that may exist between them in Figure 7. These are shown using 2D contour plots and 2 × 2 covariance matrix displayed as a curve for each pair of material properties. The gray and white matter were segmented based on their MD and FA values from DTI. The horizontal peanut shaped covariance glyph indicates large variations in shear modulus for small changes in MD. Mechanical anisotropy showed large variations compared to diffusion anisotropy. The covariance plots did not exhibit preferred a orientation indicating negligible correlation between diffusion and mechanical properties at the cardiac frequency. The gray-white matter contrast in mean diffusivity and shear modulus is investigated by plotting their 1D distributions separately (also shown in the figure). The MD distribution of gray and white matter were very similar while shear modulus distribution indicated white matter being overall stiffer than gray matter.

**Figure 7:**
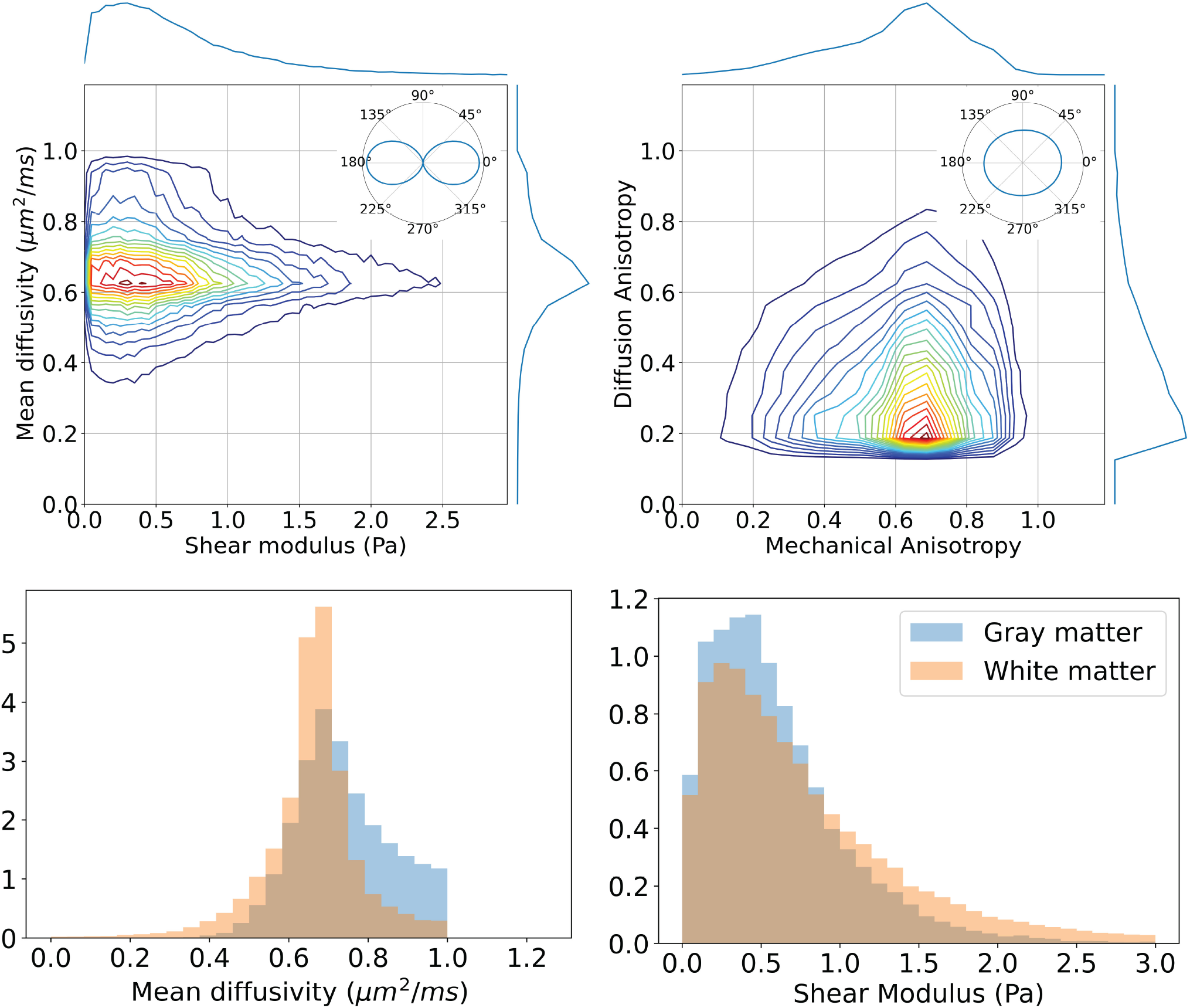
Measured correlations between diffusion and mechanical properties of brain parenchyma of a representative subject. 2D contour plots show the distribution of diffusion and mechanical properties of interest along with their 1D distributions projected on the x and y axes, and a covariance plot in the top right corner of each figure. The correlations considered are between MD vs shear modulus in both white and gray matter, and fractional anisotropy (FA) vs mechanical anisotropy (MA) for white matter only as gray matter has zero anisotropy. The individual gray-white matter distributions of mean diffusivity and shear modulus were also provided. The variation in shear modulus is much larger than that of MD as shown in the covariance plot. The results also indicate there is no significant correlation between diffusion and mechanical properties of brain parenchyma with white matter stiffer than gray matter.

We show elasticity and diffusion tensor glyphs in an axial slice from a representative healthy volunteer in Figure 8. Corpus callosum and a crossing fiber regions of interest (ROI) were chosen for illustration. The glyphs include the DT-ODF, axial, and shear stiffness tensors, which were overlaid on a FA map for anatomical reference. The stiffness glyphs pointed along the principal direction of DT-ODF due to the assumption of transverse isotropy with stiffness overall higher along the fiber direction than perpendicular to it for both the ROIs. While the DT-ODFs are homogeneous, the axial and shear stiffness glyphs were highly heterogeneous due to the different degrees of shear and axial stiffness anisotropy in each voxel.

**Figure 8:**
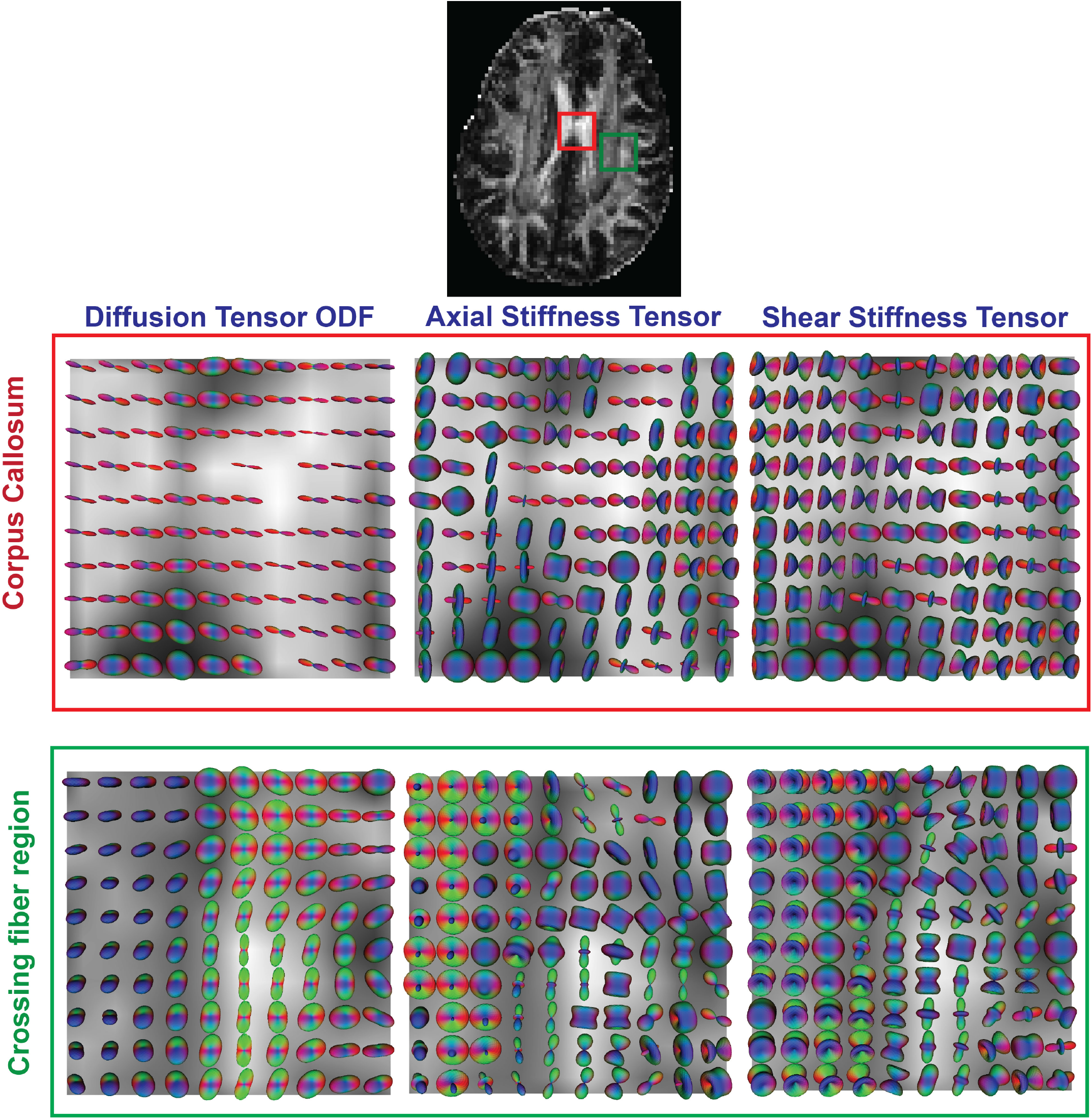
Diffusion and elasticity tensor glyphs in two regions of interest (ROI): the corpus callosum, a coherent fiber region, and the corona radiata, a crossing fiber region, in an axial slice from a representative healthy volunteer overlaid with the fractional anisotropy (FA) map for anatomical context. The glyphs include the DTI orientational distribution function (DTI-ODF), and the axial and shear stiffness tensors derived from the 4^*th*^ order elasticity tensor. While the DT-ODFs are more homogeneous, the stiffness glyphs are highly heterogeneous due to the varying degrees of axial and shear anisotropy. Notable are voxels where the principal axes of the diffusion and stiffness tensors appear parallel to one another, particularly in corpus callosum.

We show the differences in diffusion and mechanical properties across fiber tracts by overlaying them on diffusion tractograms derived from DT-ODFs for a representative subject in Figure 9. The fiber tracts were largely indistinguishable by their mean diffusivity but significant heterogeneity and anisotropy were observed in the shear modulus among fiber tracts, for example, the internal capsule, corona radiata, and U-fibers were stiffer than other white matter tracts even though they had similar MD. The stiffness was particularly pronounced in crossing fiber regions. The mechanical anisotropy was accentuated compared to diffusion anisotropy.

**Figure 9:**
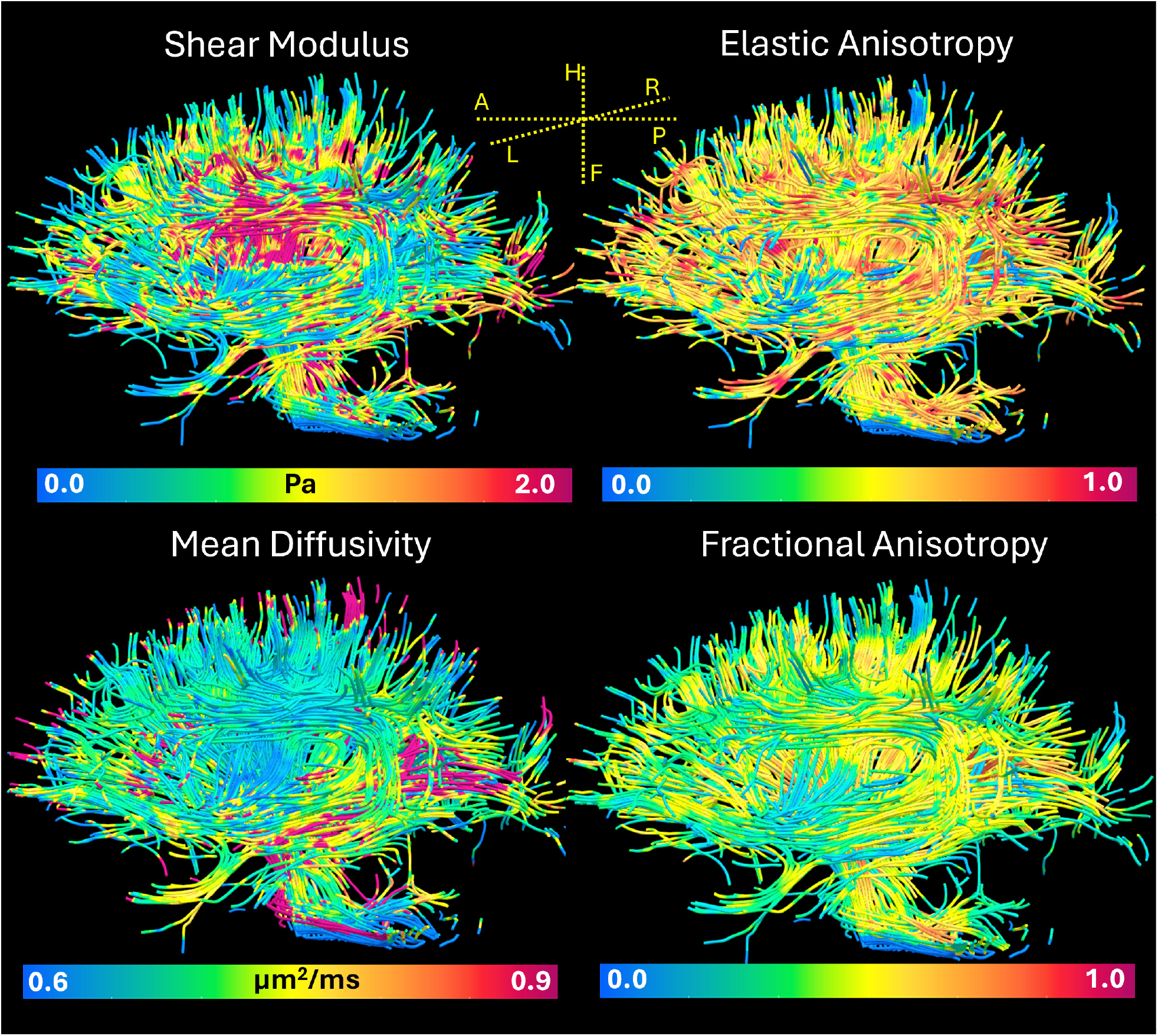
Stiffness and diffusion properties of fiber tracts in the brain shown by coloring the diffusion tractogram according to the shear modulus and mechanical anisotropy measured using MRE, and mean diffusivity and diffusion anisotropy measured using DTI. The stiffness showed significant heterogeneity across fibers with the internal capsule and corona radiata stiffer than other white matter tracts. Overall, these tracts exhibit different degrees of mechanical anisotropy which were accentuated compared to diffusion anisotropy.

Finally, we compare the results across subjects using scalar maps and tractograms in Figure 10. The shear modulus is expressed in terms of the local wavelength of the mechanical wave to adjust for heart rate differences across subjects. The results were consistent across subjects with homogeneous mean diffusivity and heterogeneous stiffness measures. Both mechanical and diffusion anisotropy values were comparable across subjects.

**Figure 10:**
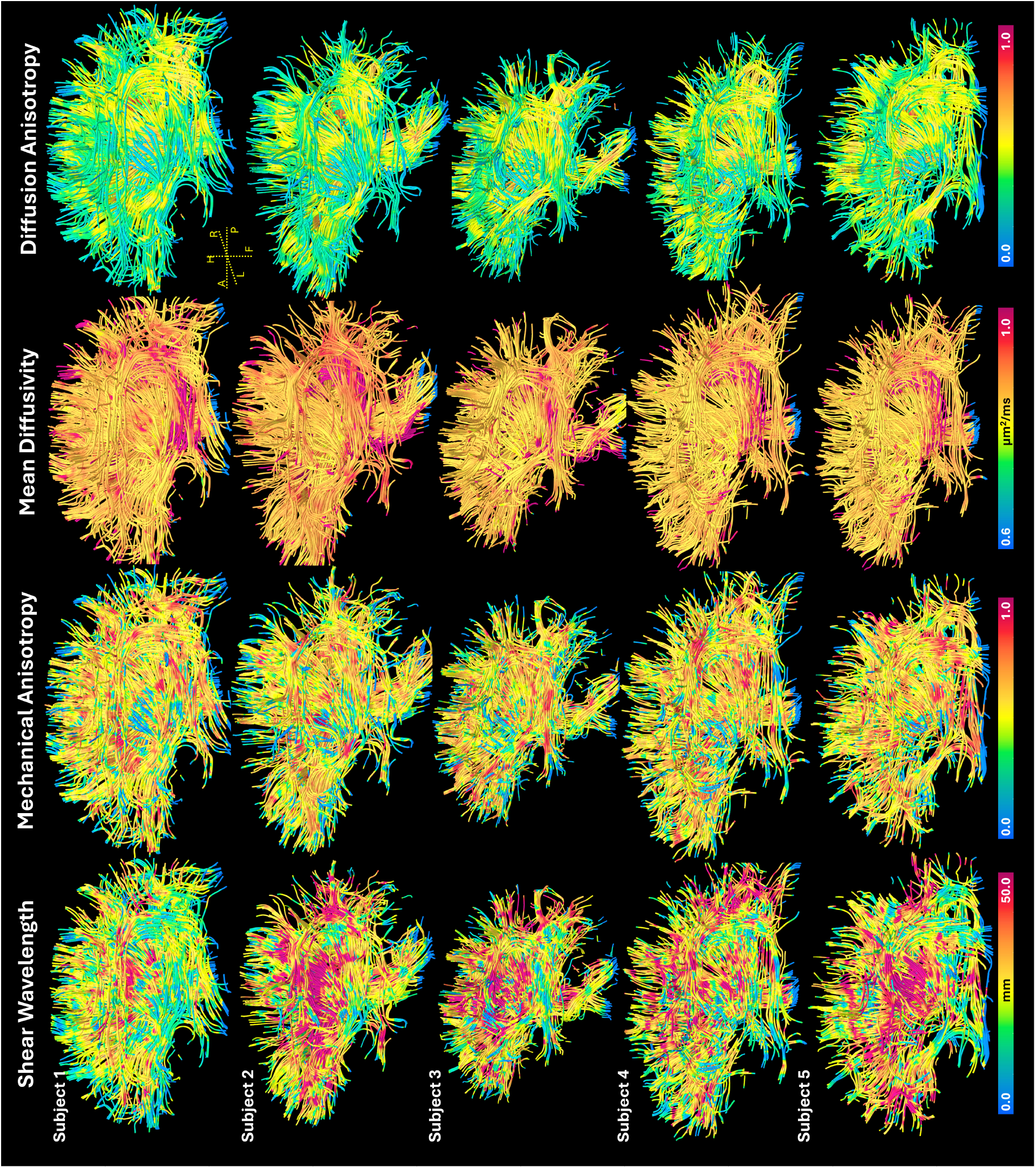
Variability in the elasticity and diffusion tensors across remaining five subjects’ brains is shown using scalar measures mapped onto fiber tracts. The stiffness measures are replaced with local wavelength to account for variations in heart rate across subjects. The maps include the mechanical anisotropy (MA), and wavelength associated with isotropic shear modulus (*G*), mean diffusivity and diffusion anisotropy. The anatomical axes (A - Anterior, P - Posterior, H - Head, F - Feet, R - Right, L - Left) are also indicated in the figure. The maps are arranged as columns and subjects as rows. The maps and the tractograms were consistent across subjects with similar values and degree of heterogeneity.

## Discussion

In this study, we present a new method to measure and map the 4^*th*^-order elasticity tensor in live human brains without requiring an external tamper or actuator. We use the principal directions of the concurrently measured diffusion tensor as *a priori* information with which to reconstruct the elasticity tensor. Our results echo previous findings but reveal new mechanical heterogeneity. We show how the diffusion measurement is affected by intrinsic brain deformation and introduce new stains and glyphs that characterize mechanical properties in each voxel. We then map mechanical properties along white matter fiber tracts illustrating their mechanical heterogeneity. We compared the results across subjects to validate our findings.

### Measuring diffusion and displacement in the pulsating brain

Measuring both intravoxel coherent and incoherent motions simultaneously in a live human brain is challenging for several reasons; 1) pulsations of the brain could affect the MR signal magnitude due to local tissue deformation [68] or from changing voxel tissue composition (i.e., time-varying partial volume effects) particularly at CSF-tissue boundaries which confound the diffusion measurement, 2) the coherent displacement induced by the heart is very small and can become conflated with incoherent diffusive displacements [69], and 3) the MRI phase that encodes coherent motion may be corrupted by gradient hardware imperfections, random head motions, etc. [32]. While cardiac gating helps mitigate some of these artifacts, it does not fully resolve them, especially in the presence of unpredictable head movements and non-linear brain deformations throughout the cardiac cycle [70]. Nonetheless, obtaining a smooth, continuous 3D displacement field is necessary for accurately reconstructing the elasticity tensor.

In this study, we have employed several strategies to overcome these aforementioned challenges. We used a standard PGSE with short gradient duration and long diffusion time to encode slow coherent motion while minimizing diffusion weighting [45]. We used image oversampling and outlier rejection with retrospective cardiac gating to correct for random MRI phase errors arising in the measurement. Furthermore, during the scanning session, we collected enough samples to retain, after outlier removal, multiple repetitions of the time-dependent complex MRI volumes at each cardiac phase. With this data we demonstrate a high degree of precision in the measurement of the displacement fields across repetitions, the observed funnel-shaped displacement profile with peak brain velocities on the order of 1 mm/s, and divergence and curl on the order of 0.2% largely agree with previous studies [32, 71]. The harmonic displacement vector plots on the three orthogonal planes show continuously varying displacement fields in 3D, free of inter-slice discontinuities also known as slice jitter [50]. The long scan time of this study is primarily due to large number of scan repetitions to obtain at least two repeats of the MRI data for each motion-encoding direction and slice for analyzing the reproducibility of the pulsations. These additional acquisitions are not needed for clinical implementation so the scan time can be reduced by at least a factor of two. Further reductions in scan time can be achieved using simultaneous multi-slice (SMS) [72], which excites multiple slices at once thereby reducing the re-binning amount.

### Effect of cardiac pulsation on DTI

We studied the effects of cardiac pulsations on DTI and correlated them with the simultaneously measured displacement fields - an analysis seldom carried out in previous studies [73, 74]. Given that the DWI signal is not just sensitive to water diffusion but to various physiological motions [75] manifesting as “pseudo-diffusion”, our proposed method has the potential to control for and possibly isolate various types of these non-diffusive motions, which can be conflated with and pose as diffusion [75], for example, affecting the MD and the apparent diffusion tensor in DTI. These physiological processes may involve cardiac and respiratory induced tissue motions, microvascular and CSF pulsations resulting in intravoxel shearing motions [68, 76], or periodic tissue compression or expansion, *inter alia*. Analyzing the complex MRI data as a function of cardiac phase, we now have a means to isolate these physiologically-induced sources of pseudo-diffusion.

Unlike the heart, which undergoes material strains as high as 20% [77], the intrinsic strains of brain tissue due to cardiac pulsations are orders of magnitude smaller (i.e., on the order of 0.2%). The resulting deformation is not sufficient to cause observable dephasing of the MR signal at the low diffusion weightings we use and does not explain the changes we observe in the DTI metrics across cardiac phases. Instead, these changes are explained by bulk tissue displacement especially at CSF-tissue boundaries, such as in the corpus callosum (CC), as evidenced by the increase in MD at certain time points in the cardiac phase. These are due to CSF in fissures above the CC being displaced into the region from head-foot piston-like brain motion, as shown in the displacement images. This was also evident in white matter fiber bundles above the ventricles that were being pushed away and pulled towards the ventricles in the H-F direction resulting in a larger FA in the ventricles when the fibers were inside and smaller when they were outside the ventricles.

### Elasticity tensor estimation

Our estimation of the elasticity tensor assumes the brain deformation at the cardiac frequency lies within the linear elastic regime. Given the measured strains are very small (i.e., “infinitesimal”), the linearity of the stress-strain relationship can be justified. However at these low actuation frequencies, the brain may be exhibiting poroelastic and not a purely elastic behavior since there maybe enough time for some fluid to move in and out of the voxel during the actuation period. This could partly explain the non-zero divergence observed by us and others [71] at the cardiac frequency. However, inverting the full poroelastic equations is not straightforward since only the average fluid and solid motions are usually observed [78]. One of the consequences of ignoring poroelasticity may be the small values of shear modulus we are observing in the brain tissue, which averages both the solid and fluid phases, given the fluid phase has zero shear modulus. The finite bulk modulus estimates may also be a result of poroelastic behavior masking the fluid flux through the voxel, given it should ideally be very large for brain tissue which is mainly composed of water. At 1 Hz excitation, it maybe expected that the shear wavelength in soft tissue becomes very long. This would imply extremely small spatial gradients of the displacement field within the tissue volume, which may make the wave equation ill-conditioned. While this may be true for some materials, we found this may not be applicable in the brain whose deformations exhibit multiple length scales. The wavenumber analysis on the measured displacement field clearly showed a continuum of wavelengths for both longitudinal and transverse waves not just the long one whose wavelength is roughly the width of the brain (100 mm). Further if the inversion is ill-conditioned, the results would be expected not to show any structure, which is not what we observed.

We applied the “cross-property” relationships arising from effective medium theory [1, 79] to make the elasticity tensor estimation problem well-conditioned. We expect the principal directions of the transverse isotropic elasticity tensor also coincide with those of the diffusion tensor in voxels with coherent anisotropic material structure [1], although their principal diffusivities and stiffness are not obviously related. These quantities have different physical units arising from different constitutive laws describing different relationships between generalized fluxes and generalized forces with governing equations potentially having different boundary and initial conditions. In the case of diffusion, the flux is a mass flux and the generalized force is a concentration gradient. In the case of the material deformation the generalized flux is the material displacement and the generalized force is the stress. These tensors are also related to and derived from different and distinct MR quantities: in DTI, from the magnitude signal for the diffusivity, and in MRE, from the phase signal. Our results comparing the diffusion and mechanical properties also show negligible correlation between the two quantities in brain tissue at cardiac frequency.

Reconstruction of the transverse isotropic elasticity tensor with *a priori* information about fiber orientation has been presented previously for muscle [27], breast [56], brain [6, 31], and phantoms [30]. Our approach differs from all these studies in several important respects, 1) we use the intrinsic pulsations of the brain instead of a tamper; 2) the low frequency of (cardiac) actuation sensitizes the signal to poroelastic effects, which may be important in normal and abnormal brain function; 3) we use standard spin echo DWI pulse sequences readily available on many scanners to measure these deformations, as opposed to DENSE [80] or other methods; 4) we obtain white matter fiber structure and orientation from a single MRI dataset instead of performing separate DTI and MRE scans, thereby eliminating registration errors due to subject motion between scans and differing eddy current distortions resulting from changes in the motion encoding gradients used in both the scans; 5) we ensure that the estimation of stiffness parameters is well-posed by introducing a physically-motivated positive definiteness constraint to improve the estimate of the five independent parameters of the transverse isotropic stiffness tensor [64]; 6) we use fast direct inversion of the displacement field using convex optimization; and 7) we introduce new glyphs and invariants for use as intrinsic quantitative imaging parameters and stains to quantify features of the elasticity tensor independent of the laboratory or patient coordinate systems.

### Heterogeneity of the diffusion and elasticity tensor fields

We see that the MD maps at different phases of the cardiac cycle are homogeneous throughout the human brain parenchyma, as was previously reported [81]. However, there is significant variability in the mechanical properties in different brain ROIs, both in gray and white matter. We ascribe these to regional differences in the meso and microscopic determinants of stiffness in complex soft matter, such as intracellular polymer content and composition, local extracellular matrix (ECM) composition and organization, stiffness variations, degree of biopolymer cross-linking and entanglement, etc. Observed discontinuities in mechanical properties at internal boundaries are expected to lead to mechanical impedance mismatches, local stress concentrations, and may predispose them to vulnerability in TBI [82]. These internal boundaries within the brain, for instance, do not appear in MD images except at CSF/parenchymal interfaces.

The higher stiffness observed in the white matter compared to the gray matter has been demonstrated in several MRE studies for a wide range of excitation frequencies [17, 24, 83], which may be due to the higher degree of crosslinking resulting from myelination [84] and higher oligodendrocyte content. The fiber tracts showed large variations in stiffness with internal capsule and corona radiata fibers exhibiting larger stiffness values compared to other fibers. This is in agreement with a study conducted on *ex vivo* human and porcine brain tissue specimens [85, 86] but opposite to that obtained on an *in vivo* MRE study at 50 Hz with a FEM analysis reconstruction [31], which maybe due to differences in the actuation frequency and analysis methods employed. Our findings could be explained by the fact that, despite having a lower axon density than corpus callosum, the corona radiata [87] is known to have higher polymer content (e.g., tubulin, actin, proteoglycans) likely due to the need to support the fanning fibers.

We do, however, note that our measured tissue stiffness values overall fall on the low end of the spectrum for brain tissue reported in the literature obtained *in vivo* and *in vitro*, which vary by orders of magnitude from Pascals (Pa) to kPa [89]. This is not surprising owing to the difficulty of making direct mechanical measurements in live brain tissue, the differences in the frequency of deformation, differences in experimental design, and the variety of mathematical/physical models used to interpret displacement, velocity, and strain data. Low values of overall average tissue stiffness, however, likely mask greater stiffness of the ECM, which occupies only about 20 percent of the tissue volume in the brain [90], the rest largely taken up by cells, which may behave like incompressible (isovolumic) micro-balloons at these timescales, with low shear stiffness [91]. Moreover, the inertial effects and extent of wave scattering are diminished while poroelastic effects can become pronounced at lower cardiac frequency, which naturally makes the material appear softer compared to that at traditionally used MRE frequencies (i.e., often around 50-60 Hz).

### Anisotropy of the diffusion and elasticity tensor fields

Diffusion anisotropy is well characterized in DTI in areas with coherent white matter organization but is known to drop in more complex white matter areas with crossing fibers, such as in parietal and frontal white matter, in deep temporal white matter, and in the centrum semiovale [81]. One could speculate that in such areas the shear stiffness could be elevated given the interdigitating fiber architecture, whereas the diffusion anisotropy would decrease there. This is evident in the elasticity tensor glyphs where the second fiber direction is accentuated in certain crossing fiber regions resulting in larger mechanical anisotropy. We would also like to note that signal noise partly contributes to the variations in both diffusion and mechanical anisotropy.

Diffusion anisotropy in gray matter is quite low, despite the higher microscopic anisotropy measured in diffusion tensor distribution (DTD) MRI studies [92–94]. There is an open question of how this microstructural motif contributes to the overall mechanical stiffness of brain parenchyma and whether we could use changes in stiffness measurements in such tissue to infer possible changes in microstructure associated with development, degeneration, diseases, and/or trauma, *inter alia*. We compared the FA from DTI to the MA (and other maps) in our MRE method. Generally, we see anisotropy appearing in white matter regions in both tensor fields, however, stiffness tensor-derived parameters tend to be much noisier compared to DTI parameters. This is partly due to differences in post-processing of the MRI data used to calculate them, with stiffness values being obtained from second numerical spatial derivatives of experimental phase parameters in MRE.

### DTI and MRE experimental designs

We are currently using the minimal number of gradient directions (i.e., six) [15] to obtain an isotropic DTI experimental design in the interest of minimizing scan time, although we recognize the value of using a greater number of gradient directions, not only to improve the estimates of the diffusion tensor in each voxel and at each phase of the cardiac cycle, but also to enable a more general higher-order tensor (HOT) representations of the diffusion and elasticity tensors. Our current design limits the material model of brain tissue we can consider to be transverse isotropic, but it is prudent to be able to employ a variety of HOT models, particularly in complex brain tissue areas, like the corona radiata. The DWI experimental design can readily be extended to accommodate High Angular Resolution Diffusion Imaging (HARDI) data [95], and methods such as mean-apparent propagator (MAP)-MRI [96], allowing us to identify complex fiber architectures in areas of the brain not adequately described by the 2^*nd*^-order DTI framework. The use of advanced gradient sampling schemes (more dense angular sampling and stronger diffusion sensitization) will also improve the accuracy of fiber tractography, especially in fiber crossing regions, and enable us to concurrently resolve and quantify the mechanical tissue properties along specific white matter pathways.

### Future challenges and opportunities

Respiratory and heart-rate fluctuations can cause some blurring and variability in the complex time-series MR signal data, and also increase noise in quantities computed from the measured displacements. We plan to improve our time-series analysis pipeline and registration methods to better account for these expected variations. This method is currently designed to be performed on subjects with fairly steady heart rates and normal blood pressure ranges, so these have to be monitored to assess data quality and improve retrospective analysis and post-processing.

Opportunities abound for applying this approach. It naturally lends itself to studying material properties of other tissues and organs in the body, particularly given recent improvements in whole-body diffusion MRI sequences, data acquisition methods and MRI hardware. Innovations in ultrahigh gradient MRI hardware should facilitate this. Innovations in diffusion MRI sequence calibration using MRI phantoms should improve the quantitative character (i.e., accuracy and precision) of both diffusivity and displacement measurements. Moreover, to obtain the MRE/DTI data, our approach does not require a custom tampers and actuators, or specially trained imaging personnel.

## Conclusion

We propose a powerful image acquisition, processing, and analysis pipeline to implement actuator-free or “intrinsic” magnetic resonance elastography (MRE) in the human brain, obtained in conjunction with diffusion tensor imaging (DTI). Besides providing estimates of bulk and shear moduli throughout the brain it also produces diffusion tensor MRI-derived quantitative “stains,” which both inform the models of mechanical properties and help provide an anatomical context, allowing us to co-register maps of brain material properties to those of anatomical features and landmarks provided by DTI. This implementation of MRE also has the potential to measure and map low-frequency material properties (both real and imaginary) and parameters in other tissues and organs throughout the body, such as the liver, providing novel “stains” and contrasts, permitting remote palpation without an external actuator or tamper.

## Supporting information

Animation 1

Animation 2

## Appendix

### Derivation of orientation averaged quantities

Let {**a, b, n**} be an orthonormal basis with **a, b** ∈ **n**^⊥^, ∥**n**∥ = 1. Any unit vector in the plane perpendicular to **n** can be written as

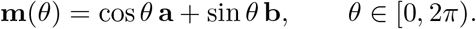

We want the uniform angular average over that circle:

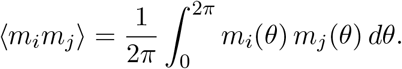

Expand the dyad:

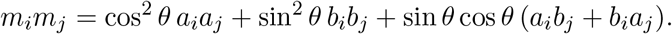

Use the elementary angular averages 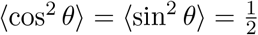, ⟨sin *θ* cos *θ*⟩ = 0, to obtain

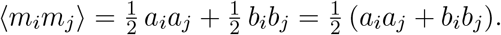

Since {**a, b, n**} is orthonormal,

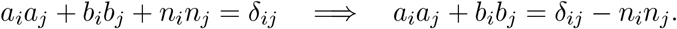

Therefore,

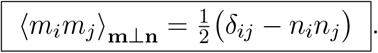

## Acknowledgments

We would like to thank Julian A. Rey for thoughtful discussions. This research was supported [in part] by the Intramural Research Program of the National Institutes of Health (NIH). The contributions of the NIH author(s) are considered Works of the United States Government. The findings and conclusions presented in this paper are those of the author(s) and do not necessarily reflect the views of the NIH or the U.S. Department of Health and Human Services. This work was also partly funded by NIH BRAIN Initiative grant 1U01EB026996-01 - “Connectome 2.0: Developing the next generation human MRI scanner for bridging studies of the micro-, meso- and macro-connectome.” and by the Military Traumatic Brain Injury Initiative (MTBI^2^) in the Department of War (DoW) under award, HU0001-22-2-0058. This work utilized computational resources of the NIH HPC Biowulf cluster (http://hpc.nih.gov). The authors have no conflicts of interest to disclose. The views, information or content, and conclusions presented do not necessarily represent the official position or policy of, nor should any official endorsement be inferred on the part of, the Uniformed Services University, the Department of War, the U.S. Government, or The Henry M. Jackson Foundation for the Advancement of Military Medicine, Inc.

## Author Contributions

KNM and PJB designed and conceived of the research project. KNM also carried out the parameter estimation, performed the experiments, and wrote the first draft of the article. AVA assisted in performing experiments and conducted the fiber tractography analysis. JES assisted in performing the clinical experiments. PJB, AVA, and JES edited the article. All authors reviewed the manuscript.

## Data availability statement

Complex MRI and physiological time series de-identified data that support the findings of this study will be made openly available in a public repository.

